# Hydrophobic organic contaminants are not linked to microplastic uptake in Baltic Sea herring

**DOI:** 10.1101/363127

**Authors:** M. Ogonowski, V. Wenman, A. Barth, E. Hamacher-Barth, S. Danielsson, E. Gorokhova

**Affiliations:** Stockholm University, Department of Environmental Science and Analytical Chemistry, Svante Arrhenius väg 8, SE-106 91, Stockholm, Sweden; Swedish University of Agricultural Sciences, Department of Aquatic Resources, Institute of Freshwater Research, Stångholmsvägen 2, SE-178 93, Drottningholm, Sweden; Stockholm University, Department of Biochemistry and Biophysics, Svante Arrhenius väg 16C, SE-106 91, Stockholm, Sweden; Swedish Museum of Natural History, Department of Environmental Science and Monitoring, P. O. Box 50 007, SE-104 05, Stockholm Sweden

**Keywords:** Microplastic, Baltic Sea, herring, hydrophobic organic contaminants, marine monitoring

## Abstract

It is commonly accepted that microplastic (MP) ingestion can lead to lower food intake and bioaccumulation of hydrophobic organic contaminants (HOCs) in aquatic organisms. However, causal links between MP and contaminant levels in biota are poorly understood and *in situ* data are very limited. Here, we investigated whether HOC concentrations in herring muscle tissue (*Clupea harengus membras*) are related to MP ingestion using fish caught along the West coast of the Baltic Sea. The MP occurrence exhibited a large geographic variability, with MP found in 22.3% of the fish examined. The population average was 1.0 MP ind^-1^; however, when only individuals containing MP were considered, the average MP burden was 4.4 MP ind^-1^. We also found that MP burden decreased with reproductive stage of the fish but increased with its body size. To predict MP abundance in fish guts, we constructed a mass-balance model using literature data on MP in the water column and physiological rates on ingestion and gut evacuation for clupeids of a similar size. The model output was in agreement with the observed values, thus supporting the validity of the results. Contaminant concentrations in the muscle tissue were unrelated to the MP levels in fish, suggesting a lack of direct links between the levels of HOCs and MP ingestion. Thus, despite their ubiquity, MP are unlikely to have a measurable impact on food intake or the total body burden of hydrophobic contaminants in Baltic herring.

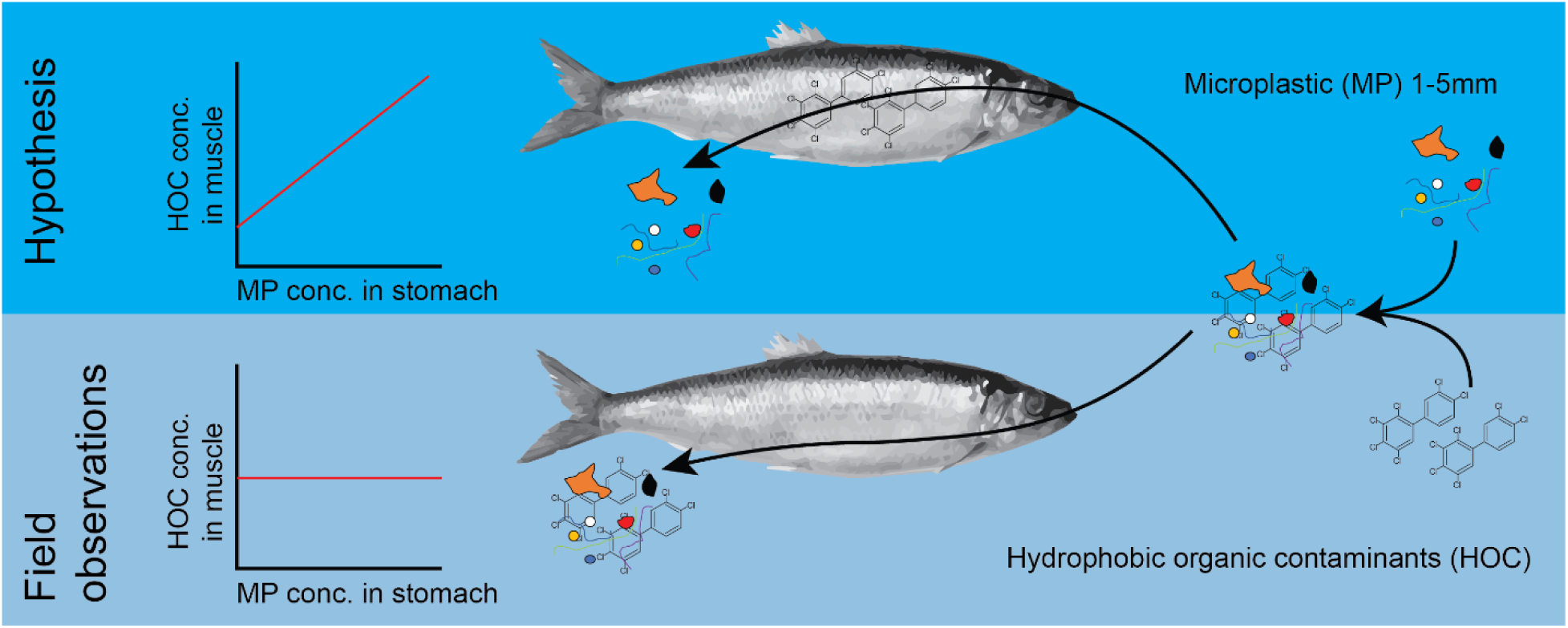

## Introduction

Plastic debris, including microplastics (MP < 5 mm), can be ingested by aquatic animals across several trophic levels (Lusher et al. 2013, 2015, Cole et al. 2013). Due to the importance of commercial fish and shellfish species for human consumption, the ingestion and presence of MP in these animals has become a matter of concern (EFSA Panel on Contaminants in the Food Chain (CONTAM) 2016). To address this concern and to provide a quantitative assessment of MP ingestion in various fish species, an active research is ongoing (Lusher et al. 2013, Foekema et al. 2013, Rummel et al. 2016, Budimir et al. 2018, Beer et al. 2018).

A commonly held paradigm states that MP ingestion can lead to decreased nutritional status (Cole et al. 2015, Ogonowski et al. 2016) and bioaccumulation of hydrophobic organic chemicals (HOCs) (Oliveira et al. 2012, Besseling et al. 2013, Rochman et al. 2013, Wardrop et al. 2016) that sorb to the MP particles in the water and desorb in the gut lumen (Mato et al. 2001, Rusina et al. 2010, Rochman et al. 2014). However, some experimental and modeling studies indicate that plastic polymers could also have a net cleaning effect acting as passive samplers while in the digestive system and thereby relieve the animals of HOCs (Gouin et al. 2011, Herzke et al. 2016, Koelmans et al. 2016, Mohamed Nor and Koelmans 2019). The relative importance of microplastics as vectors for contaminant transport remains unresolved, possibly also due to the lack of field data linking HOC concentrations in biota to ingested MP.

Here, we studied MP ingestion by Baltic Sea herring (*Clupea harengus membras* L.), a commercially exploited fish and a keystone species in the Baltic food web. Being facultative pelagic filter-feeders (Huse and Toresen 1996), herring stand a high risk of ingesting MP along with zooplankton prey and hence accumulating MP-associated contaminants. It is also a sentinel species in the Swedish National Monitoring Program for Contaminants in Marine Biota and, thus, a potential indicator species for MP monitoring in the Baltic Sea (Beer et al. 2018).

If MP ingestion indeed contributes significantly to HOC bioaccumulation in contaminated environments, then one would see a positive correlation between the amount of MP ingested over time and HOC concentrations in the herring tissues. However, there are no reliable methods to estimate accumulated MP exposure using field samples, because MP do not accumulate to any significant extent in the fish digestive system (Lusher et al. 2013, Jovanović 2017). Although gut contents reflect only a recent ingestion history (Ahlbeck et al. 2012), the MP burden determined by gut content analysis is commonly used as a reflection of the feeding habits and habitats of the fish. Another area of concern with respect to the interpretation of MP counts in environmental samples, including fish guts, is analytical accuracy and reliability of MP extraction and determination (Dehaut et al. 2016). Therefore, to increase the reliability of the MP gut content data, it is important to verify whether the recorded MP body burden is within ecologically plausible rates of ingestion and gut evacuation. To compare the observed MP abundance in the fish gut with the intake that can be expected given the MP abundance in the water column, and gut evacuation that can be expected given the food intake, a mass-balance modelling approach can be used. We applied such modeling in this study using literature-derived parameters on clupeid feeding and food processing as well as ambient MP concentrations, to estimate MP burden in the herring with a body size similar to those in our collection. We also evaluated whether HOC concentrations in the fish muscle were related to the weight-specific MP gut content of the same individual.

## Materials and Methods

### Fish collection and sample characteristics

The Baltic herring used for our analyses were collected by the Swedish National Monitoring Program for Contaminants in Marine Biota conducted by the Swedish Museum of Natural History (Stockholm, Sweden). To avoid possible bias by known point sources, we randomly selected 130 specimens that had been collected at thirteen reference monitoring stations (Figure 1), thus covering a sufficiently large geographical area that would provide a representative range of HOC and MP exposure for the analysis.

**Figure 1.**
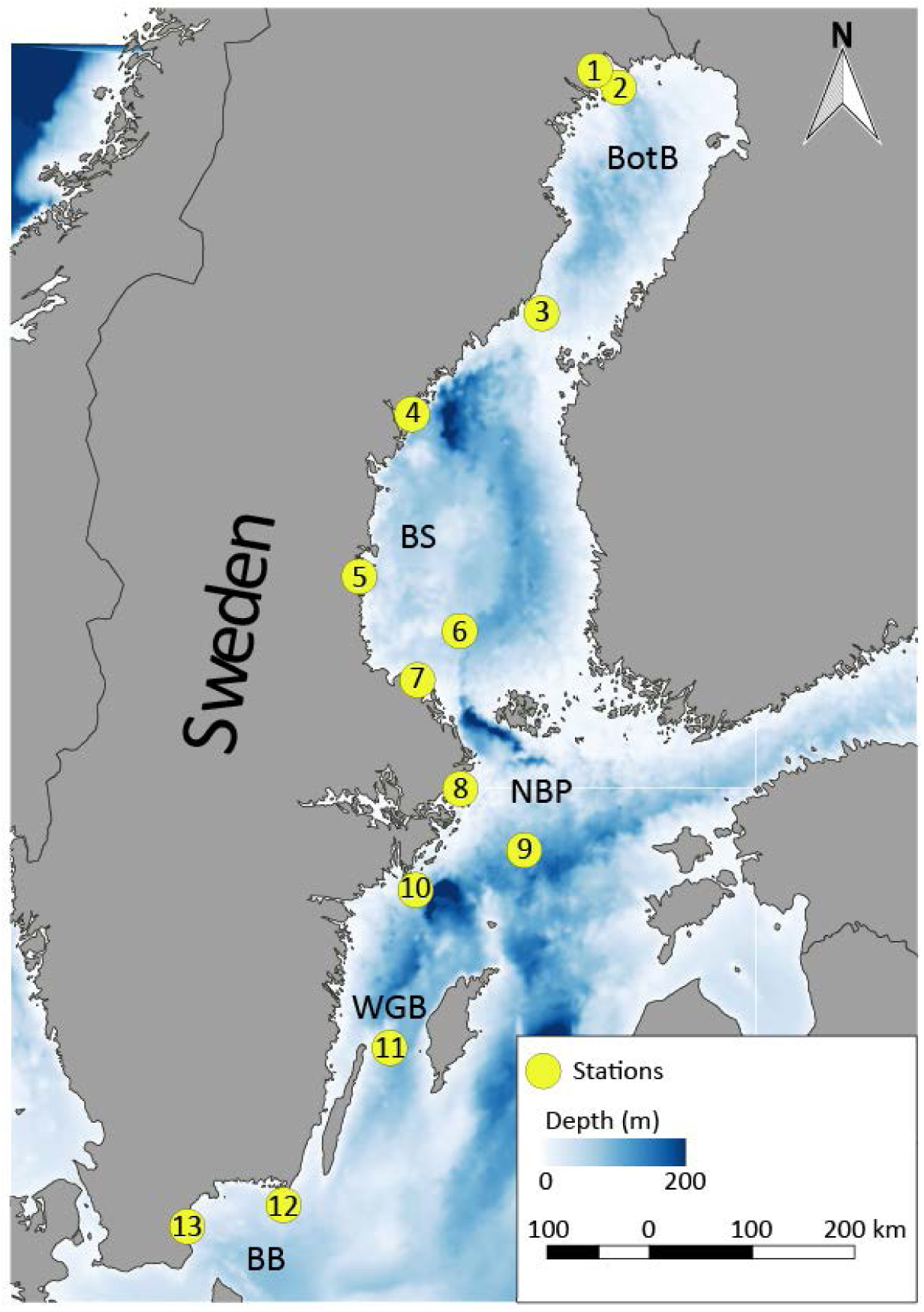
Sampling sites within the Swedish National Monitoring Program for Contaminants in Marine Biota included in this study, BotB Bothnian Bay, BS Bothnian Sea, NBP Northern Baltic Proper, WGB Western Gotland Basin and BB Bornholm Basin. 1 Rånefjärden, 2 Harufjärden, 3 Holmöarna, 4 Gaviksfjärden, 5 Långvindsfjärden, 6 Bothnian Sea offshore site, 7 Ängsskärsklubb, 8 Lagnö, 9 Baltic proper offshore site, 10 Landsort, 11 Byxelkrok, 12 Utlängan and 13 Western Hanö bight.

The sex ratio of the selected fish was approximately 50:50 and uniform across sampling sites. The individuals were 3-7 years old, with a total length of 173 ± 18 mm and body weight 35 ± 12 g (mean ± SD). The reproductive phase determined by gametocytic maturity was classified on a five-degree scale according to Bucholtz et al. (2008) and included deformed gonads (stage 1), post spawned individuals (stage 2), juveniles (stage 3), individuals with developing gonads (stage 4) and fish with mature gonads (stage 5). Each fish was dissected, and the muscle tissues taken from the middle dorsal muscle layer were used for HOC analysis, whereas the entire gastrointestinal tract (GIT) was used for the MP analysis. All sampling was performed according to standard procedures (TemaNord 1995). After dissecting, each individual GIT was packed in aluminum foil to avoid cross-contamination. All GIT samples were immediately frozen at −20 °C and stored until MP analysis at the Department of Environmental Science and Analytical Chemistry, Stockholm University, Sweden.

### MP quantification in the gastrointestinal tract of fish

Each GIT was placed in a glass Petri dish, opened with surgical scissors, and rinsed with deionized, particle-free water. Using a stereo microscope, the bolus was examined, and any items resembling MP were extracted by stainless steel pincers and transferred to clean Eppendorf tubes. The appearance of the putative MP was recorded and each particle was categorized according to its shape (fiber or fragment) and color. Hereafter, the number of MP per individual fish is referred to as *MP burden*.

To relate HOC concentration in the muscle tissue and MP intake by fish, the fish size must be taken into account when expressing MP counts. Moreover, in our collection, the bolus size varied considerably among the individual fish and geographical areas (Supporting Information Table S 2), indicating variability in feeding activity shortly before sampling and/or gut evacuation that might have been related to stress during the sampling. To account for this variability and the corresponding variation in the observed MP burden, we normalized the individual MP counts to its gut fullness; the latter was assessed by visual observation on a five-step semi-quantitative scale: 0 (empty gut, no food items), 0.25, 0.5 0.75 or 1 (full gut). The obtained values were further normalized to the individual body weight and termed *weight-specific MP burden* [number of MP / (gut fullness × body weight) (g wet weight)]. This allowed for relating HOC concentrations in the fish to the expected MP burden in the GIT on a weight basis.

The following polymer identification scheme was applied. First, to identify whether the putative MP were synthetic polymers, we followed the recommendations of Norén (2007) and Hidalgo-Ruz et al. (2012). Particles 1-5 mm in diameter were recorded and classified as MP, if all the following criteria were met: (i) uniform, unnaturally bright or of an unnatural color, (ii) lack of organic structures, and (iii) uniform diameter over the entire length of a fiber. Second, to test the accuracy of the visual identification, a random subset of 16 samples containing putative MP (i.e., the gut contents of 16 individual fish, containing in total 26 microparticles) was analyzed using Fourier transform infrared spectroscopy (FTIR). Third, MP-validation was performed by comparing the sample spectra to a published reference database (Primpke et al. 2018). The Hit Quality Index (HQI) was used to determine whether a sample spectrum matched any spectra in the database. The HQI-threshold for a match was set at 70% similarity (Thompson et al. 2004). A detailed description of the data preparation of sample spectra is provided in Supporting Information 1.1.

### FTIR analysis

The infrared spectra were recorded at 4 cm^-1^ resolution with a Bruker Vertex 70 FTIR instrument that was equipped with a Bruker Platinum attenuated reflection (ATR) unit. Data were recorded on both sides of the center burst of the interferogram during forward and backward movement of the interferometer mirror. A zero filling factor of 2 was used and the spectra were apodized with a Blackman-Harris 3-term function. The spectra for samples 1 – 6 (Supporting Information Table S 1) were recorded using a HgCdTe detector, 100 sample scans were recorded and the scanning time was 22 sec. The spectra for samples 7 – 26 (Supporting Information Table S 1) were recorded using a DTGS detector, 364 sample scans were recorded within 300 s scanning time. Individual particles were placed on the diamond crystal of the ATR unit and pressed onto the crystal with a piston. Prior to each measurement the crystal was cleaned with 99% ethanol.

### Controls and blanks

To prevent contamination by airborne particles during the examination, the dissections were performed under a Fumex local extractor (Wesch et al. 2016); each sample being analyzed for 10 min. A Petri dish filled with filtered deionized water was placed next to a test sample to serve as a blank for the quantification and characterization of potential contamination during the analysis. When working with samples, a cotton lab coat and gloves were used: moreover, the type and color of clothing were recorded to enable contamination back-tracing. All procedural blanks contained particles (mainly single fibers) of unknown origin. However, all these particles were < 1 mm and thus did not contribute to the MP counts used in the statistical analysis. If quantifiable amounts of blank contamination with particles > 1 mm were to be found, such samples would be excluded from any further analyses.

### Chemical analysis

Following the guidelines of the Swedish National Monitoring Program for Contaminants in Marine Biota, the muscle samples were analyzed for polychlorinated biphenyls (PCB 28, 52, 101, 118, 138, 153 and 180), organochlorine pesticides (DDE, DDD, DDT, HCB, AHCH, BHCH, and Lindane) and polybrominated flame retardants (BDE 28, 47, 99, 100, 153, 154 and HBCD). For most compounds, 10 g of muscle tissue from individual fish were used, whereas 1 g samples of muscle tissue from 10 individuals were pooled for a few analytically challenging compounds. An overview of the analyzed contaminants and their average concentrations in herring muscle tissue are provided in Table 1, while details of the analytical procedures and quality assurance are provided elsewhere (Bignert et al. 2016).

**Table 1.**
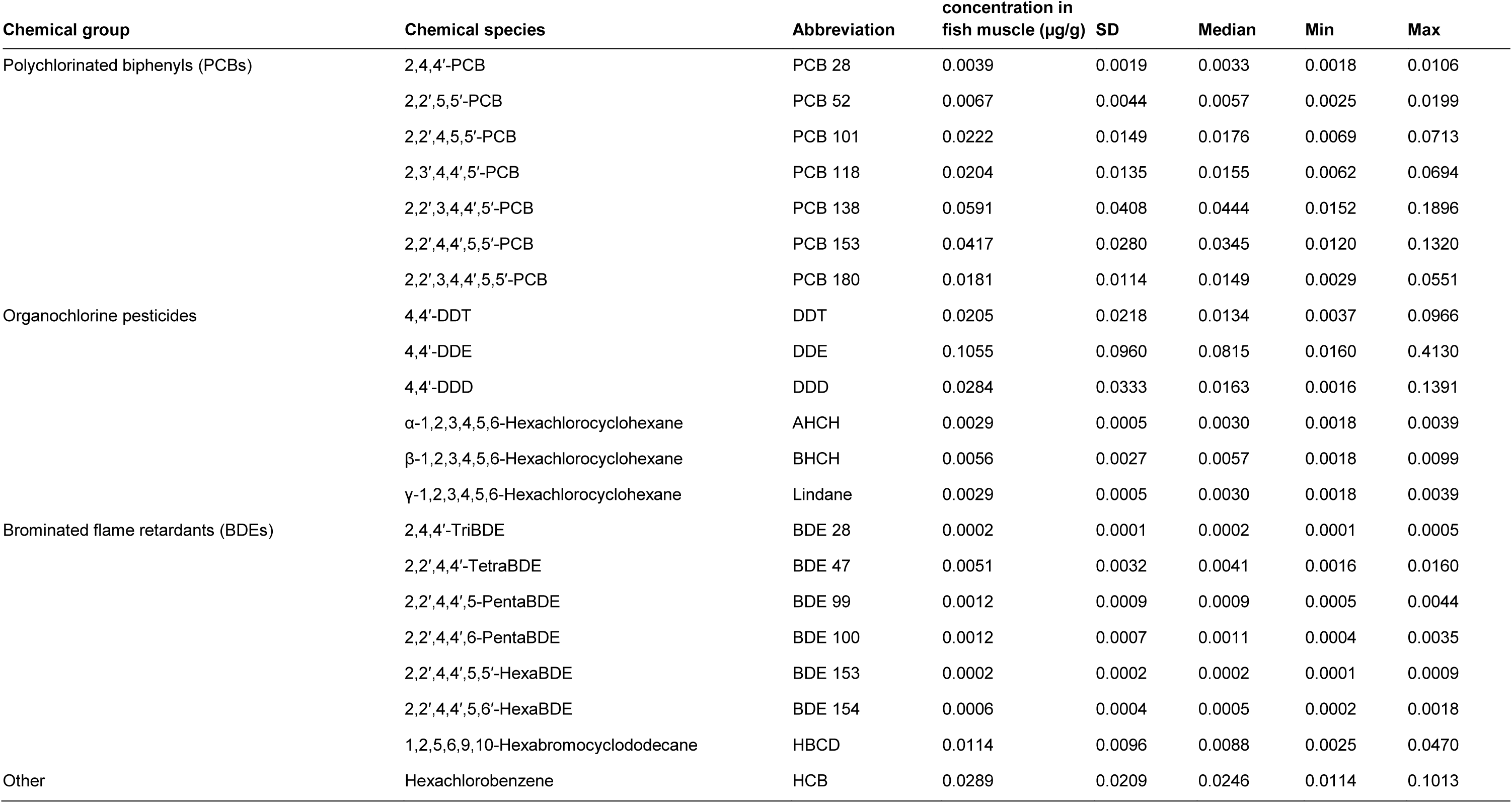
Overview of the HOCs in herring muscle tissue and descriptive statistics of their concentrations (µg g^-1^ fish muscle). SD – standard deviation.

| Chemical group | Chemical species | Abbreviation | Mean<br>concentration in<br>fish muscle ($\mu\text{g/g}$) | SD | Median | Min | Max |
| --- | --- | --- | --- | --- | --- | --- | --- |
| Polychlorinated biphenyls (PCBs) | 2,4,4'-PCB | PCB 28 | 0.0039 | 0.0019 | 0.0033 | 0.0018 | 0.0106 |
|  | 2,2',5,5'-PCB | PCB 52 | 0.0067 | 0.0044 | 0.0057 | 0.0025 | 0.0199 |
|  | 2,2',4,5,5'-PCB | PCB 101 | 0.0222 | 0.0149 | 0.0176 | 0.0069 | 0.0713 |
|  | 2,3',4,4',5'-PCB | PCB 118 | 0.0204 | 0.0135 | 0.0155 | 0.0062 | 0.0694 |
|  | 2,2',3,4,4',5'-PCB | PCB 138 | 0.0591 | 0.0408 | 0.0444 | 0.0152 | 0.1896 |
|  | 2,2',4,4',5,5'-PCB | PCB 153 | 0.0417 | 0.0280 | 0.0345 | 0.0120 | 0.1320 |
|  | 2,2',3,4,4',5,5'-PCB | PCB 180 | 0.0181 | 0.0114 | 0.0149 | 0.0029 | 0.0551 |
| Organochlorine pesticides | 4,4'-DDT | DDT | 0.0205 | 0.0218 | 0.0134 | 0.0037 | 0.0966 |
|  | 4,4'-DDE | DDE | 0.1055 | 0.0960 | 0.0815 | 0.0160 | 0.4130 |
|  | 4,4'-DDD | DDD | 0.0284 | 0.0333 | 0.0163 | 0.0016 | 0.1391 |
| | α-1,2,3,4,5,6-Hexachlorocyclohexane | AHCH | 0.0029 | 0.0005 | 0.0030 | 0.0018 | 0.0039 |
| | β-1,2,3,4,5,6-Hexachlorocyclohexane | BHCH | 0.0056 | 0.0027 | 0.0057 | 0.0018 | 0.0099 |
| | γ-1,2,3,4,5,6-Hexachlorocyclohexane | Lindane | 0.0029 | 0.0005 | 0.0030 | 0.0018 | 0.0039 |
| Brominated flame retardants (BDEs) | 2,4,4'-TriBDE | BDE 28 | 0.0002 | 0.0001 | 0.0002 | 0.0001 | 0.0005 |
|  | 2,2',4,4'-TetraBDE | BDE 47 | 0.0051 | 0.0032 | 0.0041 | 0.0016 | 0.0160 |
|  | 2,2',4,4',5-PentaBDE | BDE 99 | 0.0012 | 0.0009 | 0.0009 | 0.0005 | 0.0044 |
|  | 2,2',4,4',6-PentaBDE | BDE 100 | 0.0012 | 0.0007 | 0.0011 | 0.0004 | 0.0035 |
|  | 2,2',4,4',5,5'-HexaBDE | BDE 153 | 0.0002 | 0.0002 | 0.0002 | 0.0001 | 0.0009 |
|  | 2,2',4,4',5,6'-HexaBDE | BDE 154 | 0.0006 | 0.0004 | 0.0005 | 0.0002 | 0.0018 |
|  | 1,2,5,6,9,10-Hexabromocyclododecane | HBCD | 0.0114 | 0.0096 | 0.0088 | 0.0025 | 0.0470 |
| Other | Hexachlorobenzene | HCB | 0.0289 | 0.0209 | 0.0246 | 0.0114 | 0.1013 |

### Data analysis and statistics

#### Relationships between biological factors, geography and ingested microplastic

We used generalized additive models (GAM) in package *mgcv* to examine relationships between specific biological variables (*weight*, *gut fullness*, *age* and *reproductive phase*) and *MP burden*; *sea basin* was used as a random factor in the model since the inclusion of this term lowered the Akaike Information Criterion from 323 to 270. The model was specified as:

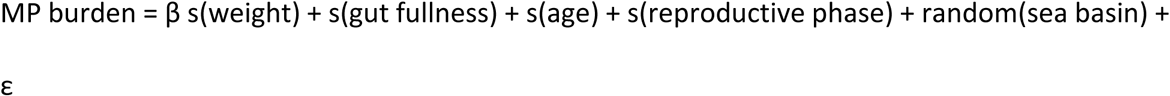

The multicollinearity between the explanatory variables was evaluated as low (< 0.38) using concurvity measures (Amodio et al. 2014) calculated by the *mgcv*-package. Due to the overrepresentation of zeros in the data (overdispersion) for the MP burden, the model was run using zero-inflated Poisson error structures. Model performance was assessed using residual plots. Differences in the MP burden between the basins were tested using Permanova with *station* nested within *basin* as a random factor (Anderson 2001). The significance level was set at α = 0.05; all statistical analyses were conducted in R 3.5.0 (R Core Team 2014).

#### Relationships between HOCs and ingested microplastic

Maximum-likelihood Factor Analysis with Varimax rotation was used to assess the degree of association between the chemical variables and weight-specific MP burden in the GIT. Prior to the analysis, Bartlett’s test of sphericity was performed to confirm patterned relationships between the variables and was statistically significant (χ^2^_15_ = 176, p <0.0001). A scree plot was used to determine the number of factors to retain, and factor loadings > 0.7 were considered statistically significant (MacCallum et al. 2001). When measured values were below the limit of quantification (LOQ), they were imputed by LOQ divided by the square root of two (Succop et al. 2004). The analyzed chemical concentrations were summed and grouped into their respective contaminant groups (PCBs, PBDEs and organochlorine pesticides).

#### Modeling plastic ingestion by herring

To evaluate whether the observed MP burden could be predicted using ambient MP abundance data and food processing rates, we modeled the ingestion of MP using literature-derived parameters on food uptake, egestion, and MP abundance in the study area. The rationale is that observed MP abundance in the gut would reflect average exposure levels assuming that (1) MP concentrations are fairly homogeneous in the outer coastal areas (Gorokhova 2015, Gewert et al. 2017), which are the main feeding grounds of herring (Flinkman et al. 1998), (2) the MP abundance in the water column, where the fish feed, is similar to that at the surface, where the data on the relevant size fraction of MP (1-5 mm) were collected; (3) MP ingestion by herring is non-selective and thus proportional to the MP abundance in the water, and (4) gut evacuation rates are non-discriminatory, i.e., MP are egested at the same rate as prey remains. Then, the MP burden (MP ind^-1^) at any given time, *t*, can be written as the mass balance between the uptake and loss rates (Eq. 1):

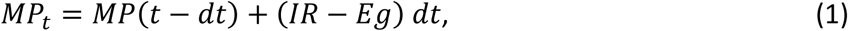

where *IR* and *ER* are the ingestion and egestion rates (MP h^-1^), respectively. They can be calculated as:

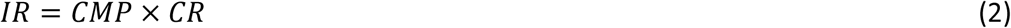

and

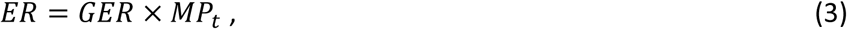

where *CMP* is the ambient MP concentration (number of MP L^-1^), *CR* is the clearance rate (L h^-1^; the volume of water swept clear of particles per individual and hour), and *GER* is the gut evacuation rate (h^-1^).

We used literature data to parameterize the model (Supporting Information Figure S 1, Table S 3). The MP concentrations in the target size range (1-5 mm) from surface waters in the outer Stockholm archipelago (Gewert et al. 2017) were used as CMP values. Clearance rates were estimated using reported feeding rates for North Sea herring on *Calanus finmarchicus,* a copepod of similar size as the microplastics considered here, and the main prey for herring (Varpe and Fiksen 2010) (see Supporting Information 2.1 for the calculation of CR). As published gut evacuation rates for adult herring were not available, we used experimental values reported for other clupeids of similar size, European pilchard (*Sardina pilchardus*) (Costalago and Palomera 2014) and South American pilchard (*Sardinops sagax*) (van der Lingen 1998), which have similar feeding ecology and physiology as Baltic herring (Collard et al. 2017). The physiological rates used in the model corresponded to the average size of our fish.

The model was implemented using STELLA^®^ ver. 9.4.1 software (iSee systems, Inc. Lebanon, NH, U.S.A.) to estimate MP burden (MP ind^-1^) dynamics in a fish population at a given MP abundance. The intrapopulation variability was simulated using a Monte Carlo generator with 1000 permutations (details on the simulation settings are provided in Supporting Information 2.2). To validate the model, we compared the simulated data distribution from the model to the field data using descriptive statistics, χ^2^, and the two-sample Cramér-von Mises tests.

## Results

### Observed MP burden

Particles identified by visual inspection as MP were found in 44 out of the 130 individuals (33.8%; range: 0 to 51 pieces of plastic fiber or fragments ind^-1^). In these 44 individuals, the mean abundance was 7.8 ± 12.2 particles ind^-1^ (± SD). The dominant type of the MP were fibers of various colors (87.6%), while fragments were less frequent (12.4%). However, only 38.5 % of the putative MP were classified as either synthetic or semi-synthetic (viscose) by FTIR and 38.5% were classified as being of natural origin (e.g. chitin, fur and cellulose); 23 % could not be identified (Supporting information Table S 1). None of the samples exclusively matched the most commonly found polymers in the environment; polyethylene (PE), polypropylene (PP), polystyrene (PS), polyethylene terephthalate (PET) and polyvinylchloride (PVC).

After correcting for the proportion of the misclassified samples, only 17 individuals containing MP remained (13.1 %; range: 0 to 20 MP), with a mean abundance of 4.5 MP ind^-1^ ± 5.3. When all examined individuals were considered, the population average was 0.9 MP ind^-1^, with the 95% bootstrap confidence interval ranging 0.5 - 1.6 MP ind^-1^. The variation in the MP burden between the stations and basins was high (Figure 2, Supporting information Table S 2) and no significant differences in the MP burden between the basins were found (*station* nested within *basin* as a random factor, pseudo F_4,117_ = 0.9, p = 0.49).

**Figure 2.**
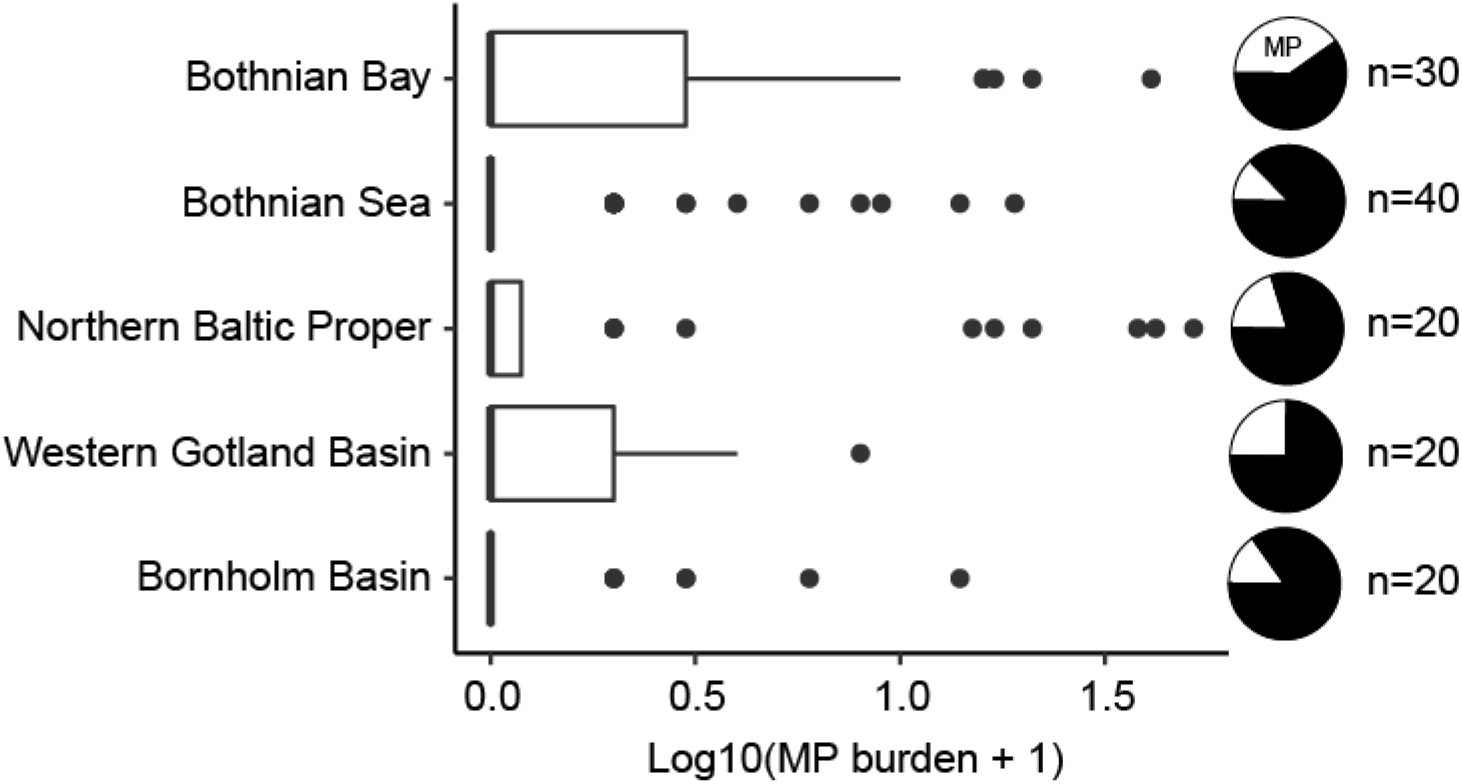
Boxplot of Log10-transformed MP abundance in the gastrointestinal tract (GIT) of herring per basin ordered from north to south. Data are presented as medians (vertical lines), inter quartile range, IQR (boxes), 1.5 IQR (whiskers) and outliers (points) being > 1.5 IQR. The black slices of the pie charts indicate the proportion of examined herring with no MP in the GIT.

### Predicted vs. observed MP burden and frequency of occurrence

The model predicted that 81% of fish contained MP, with a mean MP burden of 4.7 MP ind^-1^; these values were about five times as high as the observed values. The ranges of the frequency distributions for the simulated and observed values were overlapping, although the field observations were more strongly skewed towards zero values compared to the model prediction (Figure 3 A; Supporting Information Table S 4). The difference between the distributions was statistically significant (Cramér-von Mises T = 186, p < 0.0001). However, when zero values were excluded, the distributions, albeit still significantly different (Cramér-von Mises T = 10.6, p < 0.01), became more similar (Figure 3 B; Supporting Information Table S 4), indicating that much of the difference between the distributions was driven by the significantly higher proportion of zero observations in the field data (χ^2^ = 219.5, p < 0.0001).

**Figure 3.**
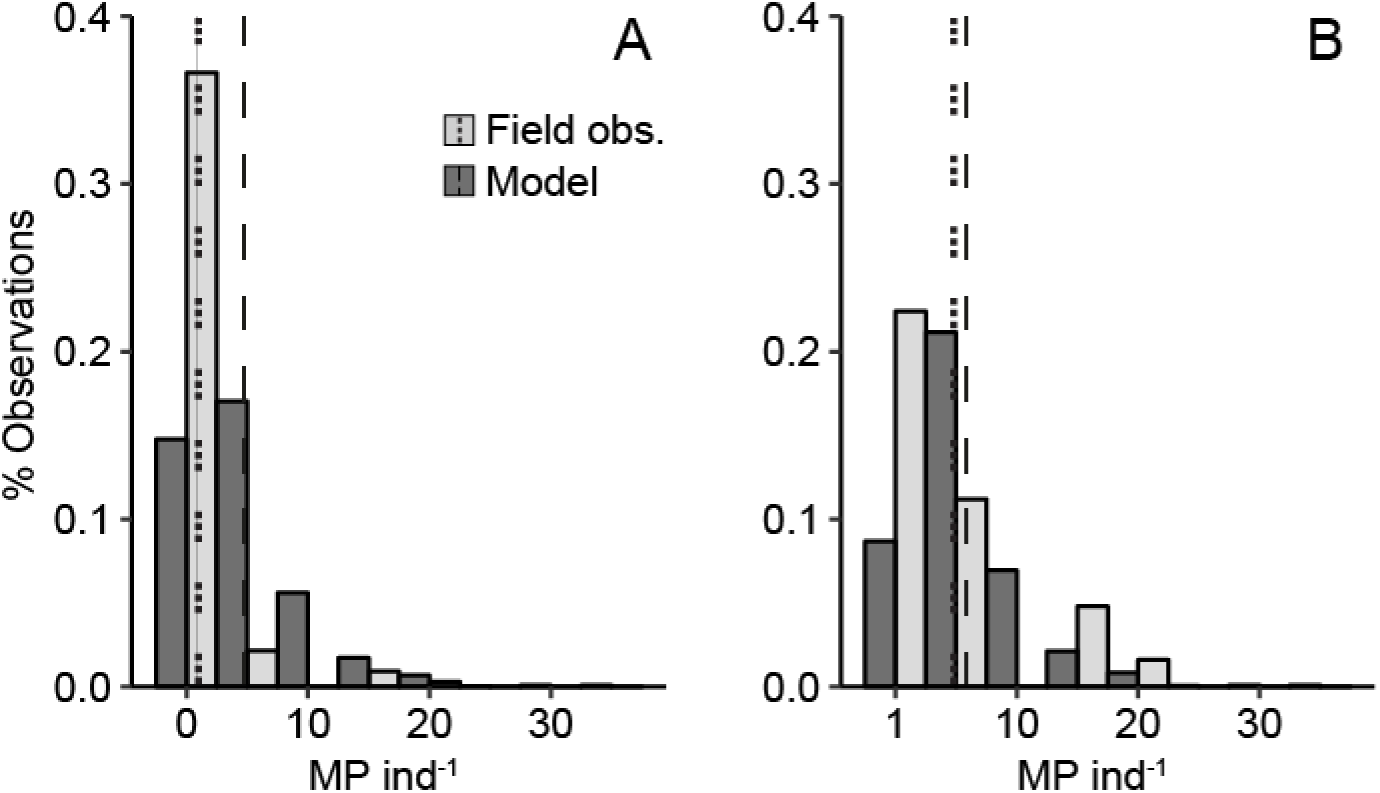
Frequency distribution of the MP burden based on the model simulations (dark grey bars) and field observations (light grey bars). Panel A shows the entire dataset and panel B presents only fish with MP in the GIT (i.e., the non-zero values). The dashed vertical lines indicate the mean values for the model simulations (long dash) and the observations (short dash).

### Linkage between MP intake and HOCs

We found no relationship between the weight-specific MP burden and the concentration of any of the HOCs (Figure 4). Together, the two factors explained a cumulative variance of 84.4% (Supporting Information Table S 5). As a variable, weight-specific MP burden loaded weakly and negatively (−0.14) on the first axis and moderately positive (0.58) on the second axis. In contrast, the organochlorine pesticides and PBDEs loaded significantly and positively on the first axis, while the PCBs loaded moderately positive (0.56) on the first and significantly positive (0.82) on the second axis. Hence, no contaminant group had loadings clustering with those for the weight-specific MP burden.

**Figure 4.**
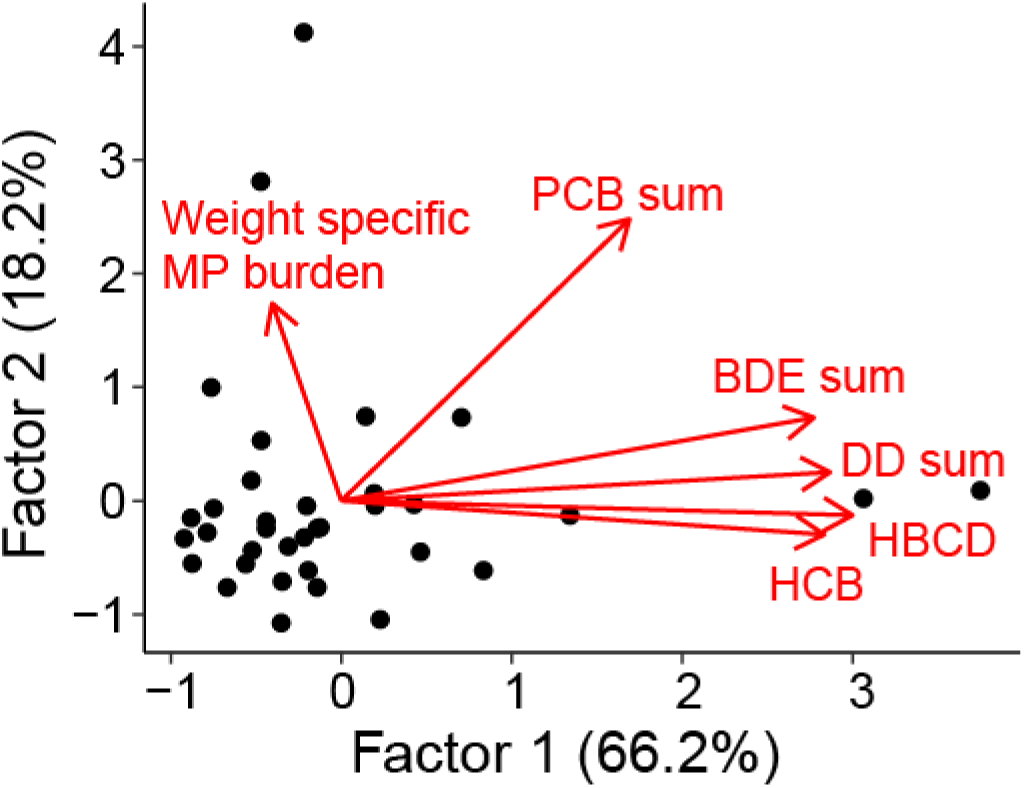
Factor scores (axes) and loadings (arrows) of contaminants (HBCD, HCB and the sum of PCBs, BDEs and DDs) and weight-specific MP burden.

### Biological factors related to MP burden

The MP burden was positively and nearly linearly related to fish *body weight* (GAM, χ^2^ = 13.1, p < 0.01, Figure 5 A). In contrast, a negative effect was found for *reproductive phase*, where MP burden was significantly lower in fish that had reached sexual maturity (GAM χ^2^ = 16.4, p < 0.01, Figure 5 B). *Gut fullness* only had a negative effect on MP burden when the GIT was empty of food items (GAM χ^2^ =39.4, p < 0.0001, Figure 5 C) while *Age* displayed a weak negative relationships with MP burden (GAM χ^2^ = 16.6, p < 0.001, Figure 5 D).

**Figure 5.**
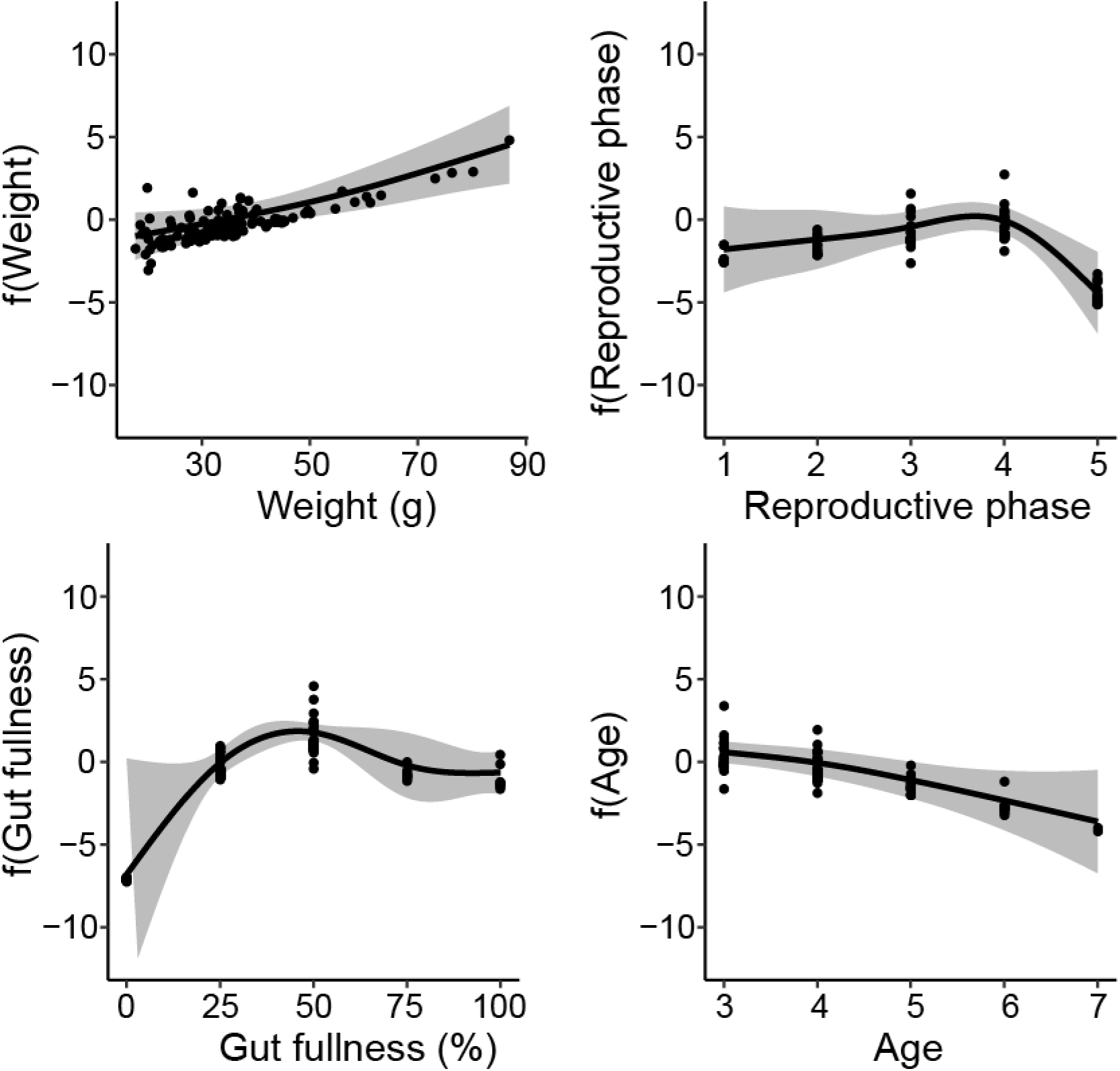
Generalized additive models (GAMs) showing partial response curves for the explanatory biological variables: *body weight* (A), *reproductive phase* (B), *gut fullness* (C) and *age* (D). The classes for *reproductive phase* correspond to: 1 = deformed gonads, 2 = post spawned, 3 = juvenile, 4 = developing gonads and 5 = mature gonads. The vertical axis shows the relative influence of the explanatory variable on the prediction of MP burden on the base of partial residuals. Grey bands indicate 95% confidence interval for each curve.

## Discussion

### Microplastics are common but not abundant in herring guts

Microplastics (mostly fibers of synthetic and semi-synthethic origin) were found in about 20% of the fish. While these values are in good agreement with those reported for herring by Beer et al. (2018) for the central Baltic Sea (20% containing MP, with 93% fibers), other studies report considerably lower MP frequency of occurrence and fiber contribution to total MP in herring. Both Foekema et al. (2013) and Rummel et al. (2016) found plastics in only 2% of herring samples from the North Sea and the Southern Baltic Sea, with fibers accounting for less than 10% of MP. Having excluded fibers from their analyses, Budimir et al. (2018) reported a frequency of occurrence as low as 1.8% in herring from the northern Baltic Sea. These discrepancies between different studies could be related to differences in fish size and gut fullness. For example, Foekema et al. (2013) used fish that were considerably larger (>200 mm total length) which most likely already had switched from filter feeding to raptorial feeding on larger prey (Huse and Toresen 1996). This change in feeding mode would result in a lower ingestion rate of zooplankton-sized plastic particles and thus in a lower overall MP burden. In the study of Rummel et al. (2016), many fish stomachs were empty, which probably was related to arrested feeding in concert with spawning, and, possibly, stress-induced gut evacuation caused by the fish sampling (Wilkins 1967, Vinson and Angradi 2011). This lends further support to our findings that MP burden increases with fish size (Beer et al. 2018) and decreases with reproductive phase (Stacey and Hourston 1982). In addition, one would expect the amount of ingested MP to scale with the absolute size of bolus or gut fullness. However, since this relationship was weak (Figure 5 C), our findings only partly support this expectation. One possible explanation for this could be slower egestion of MP compared to prey, similar to the selective retention of plastic fibers in amphipods (Au et al. 2015) and fragments in cladocerans (Ogonowski et al. 2016), which would result in a temporary accumulation of MP in the fish gut and obscure the expected positive relationship between the gut fullness and MP burden. While fish size appears to be the strongest covariable for standardizing gut MP content, gut fullness was also influential, particularly for fish with empty guts, which may occur during fasting periods (Darbyson et al. 2003). Although the effect of *Age* also was statistically significant, the effect was not particularly strong and most probably of low biological importance.

The range of the MP burden predicted by our simple model was similar to that observed in the field caught specimens, although the proportion of fish predicted to contain MP was more than fivefold higher (Figure 3 A). This is, however, not surprising because, the frequency of zero values was driven by the variability in MP occurrence in the water that was derived from surface-collected MP. Moreover, the model assumed homogeneous MP distribution in the water column, which is unlikely, because the MP distribution is patchy varying with depth (Gorokhova 2015). Also, MP can form aggregates that are too large to be mistaken for food (Long et al. 2015, Lagarde et al. 2016). Therefore, the distribution of MP concentrations originating from surface collections and used to model MP encounter rate might not reflect the actual abundance of MP available to the fish. The observed MP burden for the population was also more variable, which is likely to be related to diel variations in feeding and gut evacuation under natural conditions (Seyhan and Grove 2003), not accounted for by the model. Other biological factors, such as maturity level, ontogenetic changes in feeding, and behavior, may have affected the probability of MP ingestion and thus contributed to the intrapopulation variability in the MP burden. Finally, fishing methods (which may induce gut evacuation) and time of capture (which may reflect diurnal differences in feeding activity) may have contributed to the observed discrepancy in the MP burden distribution. Nevertheless, given the simplicity of the model and the uncertainties associated with its parameters, the predicted values were sufficiently close to those found in the field, indicating that MP uptake can be predicted provided that we have reliable MP abundance estimates.

### No correlation between weight-specific MP burden and HOCs

The transfer of hydrophobic contaminants from ingested plastics to biota has been described as the so-called “Trojan horse” effect (Cole et al. 2011). While this transfer has been demonstrated under laboratory conditions (Besseling et al. 2013, Rochman et al. 2013, Browne et al. 2013, Batel et al. 2016), recent modelling studies indicate that natural sources are much more important than MP in explaining HOC bioaccumulation patterns in aquatic organisms (Koelmans 2015, Koelmans et al. 2016, Mohamed Nor and Koelmans 2019). We did not find any correlation between HOC concentrations in herring muscle and MP burden, although it could be argued that omitting small MP (< 1 mm) from our analysis, could have biased the results. Indeed, by focusing on the larger MP, we ignored the potentially important influence of a higher total surface area and thus higher HOC desorption rates (Hendriks et al. 2001, Hartmann et al. 2017). However, the ingestion of such small particles by fish of this size is rather unlikely, because filter-feeding herring have a relatively low capacity to retain small particles due to their rather wide gill raker spacing (Gibson 1988, Collard et al. 2017) and actively avoid smaller prey while feeding raptorially (Aro et al. 1989, Casini et al. 2004). In fact, this line of reasoning has been supported by several other studies reporting a predominant retention of MP of > 1 mm by similarly-sized herring (Lenz et al. 2016, Collard et al. 2017, Beer et al. 2018). Moreover, given the short residence time (Grigorakis et al. 2017) of ingested plastics and the slow desorption kinetics of many HOCs, the lack of correlation between the MP and organic contaminants is rather expected and in line with other reports for fish and other aquatic animals (Herzke et al. 2016, Rehse et al. 2018, Kleinteich et al. 2018).

Causality is difficult to prove using environmental samples, where many different parameters may affect contaminant body burden of an organism (Hartmann et al. 2017), including various biotic factors that have significant effects on both MP (this study) and HOC levels (Persson et al. 2013, Silva Barni et al. 2014). However, our findings suggest that there is no tenable relationship between the MP intake and tissue contaminant concentrations in the Baltic herring (Figure 4). Similarly, no correlation has been found between the amount of ingested plastic and HOC concentrations in northern fulmars (*Fulmarus glacialis*) from the Norwegian coast (Herzke et al. 2016), even though the birds had ingested much larger amounts of plastic and their gut passage time for plastic debris is several orders of magnitude longer than in herring (Ryan 2015). This lack of relationship is also supported by the relatively constant MP burden observed in Baltic herring over the past three decades (Beer et al. 2018), while muscle concentrations of HOCs have decreased significantly (Bignert et al. 2016). The mass-balance model indicates that our measurements of the MP burden are ecologically plausible given the currently reported abundances of MP in the Baltic surface water, thus supporting the reliability of the MP burden estimates in the Baltic herring reported here and in other studies and providing confidence in the methods employed. Taken together, these findings contrast the currently held paradigm that microplastics are an important source of HOCs in aquatic organisms (Mato et al. 2001, Rochman et al. 2013).

## Acknowledgements

This work was supported through the Joint Programming Initiative Healthy and Productive Seas and Oceans (JPI-Oceans) WEATHER-MIC project by the Swedish Research Council for Environment, Agricultural Sciences and Spatial Planning (FORMAS) [grant number 942-2015-1866], FORMAS project; ir-PLAST [grant number 2015-932], through the joint Baltic Sea research and development programme (BONUS) MICROPOLL project by the Swedish Innovation Agency VINNOVA [grant number 2017-00979] and the Baltic Ecosystem Adaptive Management (BEAM) project. The study was designed, analyzed and compiled solely at the responsibility of the authors without the involvement of the funding agencies. Preliminary results were presented at the 15^th^ International Conference on Environmental Science and Technology in 2017, Rhodes, Greece, paper number: CEST2017_00690. A preprint of this manuscript has been deposited in BioRxiv; DOI: https://doi.org/10.1101/363127.

## Notes

**Declarations of interest**: none

