## Supplemental Information for "Hydrophobic organic contaminants are not linked to microplastic uptake in Baltic Sea herring"

### 1.1 Data handling of FTIR spectra

The original FTIR spectra contained residual signals from atmospheric $CO_2$. These were subtracted using five spectra recorded without sample in which the regions without $CO_2$ signals were replaced by straight lines. Data outside the range of 3845 to 400 $cm^{-1}$ were ignored. In order to treat the data in the same way as those in the reference data base (Primpke et al. 2018) the spectra were baseline corrected (64 baseline points, 10 iterations) with the OPUS 5.5 software. Their wavenumber values were then made compatible with those in the reference data base using one of its spectra as the principal file.

To reduce noise and enhance the spectrum quality without losing subtle spectral information, each spectrum passed through a baseline correction and denoising procedure using a second order derivative Savitzky-Golay filter to minimize baseline effects (Renner et al. 2019) as implemented in Unscrambler® X (version 10.5.1; Camo Software, 2018). Pre-processed spectra were then compared to the spectral libraries assembled by Primpke and co-workers (Primpke et al. 2018) using BioRad KnowItAll® Informatics System software. The Correlation algorithm implemented in the software was used to evaluate each query spectrum to the spectra of the databases. The most appropriate match (Hit Quality Index; HQI) was selected based on matching peak wavenumber positions and a minimum 70% correlation between unknown and matched spectra. In all spectra, we focused on the 3799 to 850 wavenumber ($cm^{-1}$) region, excluding the interval 2690 to 1900 wavenumber ($cm^{-1}$).

### 2.1 Model parameterization

#### *2.1.1 Clearance rate (CR)*

Clearance rates (CR) for Baltic herring were calculated based on intake rates of *Calanus finmarchicus* in the North Sea (Fig. 5 in Varpe and Fiksen 2010), which is the main prey for herring in this area. These copepods are also of similar size (2-3 mm prosome length) as the microplastic particles considered in this study (Pasternak et al. 2004). We used the reported values on *C. finmarchicus* consumption by herring expressed as (J copepod [J herring]$^{-1}$ day$^{-1}$) and ambient *C. finmarchicus* abundance to obtain the CR.

The average CR of the Baltic herring population examined in this study (L $h^{-1}$) was calculated assuming an energy content of 3.5 kJ and 10 kJ g wet weight$^{-1}$ for *C. finmarchicus* and herring, respectively (Varpe et al. 2005). The average weight of the sampled Baltic herring (35 g) was used to derive the consumption rate on an individual basis using a first-order exponential decay function fitted to data on the CR and prey abundance for the North Sea herring feeding on *C. finmarchicus* (Figure S 2). The asymptote value ($1.04 \times 10^3$ L ind.$^{-1}$ $h^{-1}$) was assumed to represent CR of the Baltic herring, because mesozooplankton abundance in the Baltic Sea normally supersede the maximum reported abundance for *C. finmarchicus* in the North Sea (Varpe and Fiksen 2010, Gorokhova et al. 2016).

*2.1.2 Ambient MP concentrations in the Baltic Sea (CMP)*

We used the average microplastic concentrations reported by Gewert et al. (2017) in the outer Stockholm archipelago (0.58 MP $m^{-3}$) estimated by surface manta trawls (335 µm mesh). These values were used, because the size range (median MP size and inter quartile range, IQR: fragment diameter = 1 mm (IQR 0.6-1.5 mm), fiber length = 1 mm (IQR 1-3 mm)) fits well the size of MP recovered from the fish guts. Also, the polymer materials have been rigorously identified by FT-IR in this selection of the field-collected MP, thus ensuring that the fragments collected were indeed microplastics.

*2.1.3. Gut evacuation rates (GER)*

We were not able to find data on gut evacuation rates for adult herring; therefore, a lower and an upper limit reported for two clupeid species of similar size and feeding ecology as the herring analyzed here (Collard et al. 2015) were chosen. The lower limit (0.05 $h^{-1}$) was adopted from the experimental and field data collected for adult South American pilchard (*Sardinops sagax*) (van der Lingen 1998). The upper limit (0.26 $h^{-1}$) was experimentally derived for adult European pilchard (*Sardina pilchardus*) (Costalago and Palomera 2014).

### 2.2 Monte Carlo simulation of MP burden in the Baltic herring

To estimate MP burden (MP $ind^{-1}$) dynamics at a given MP abundance from time 0 to the point when it is stabilized (48 h), we performed Monte Carlo simulation with 1000 permutations using STELLA® ver. 9.4.1 software (iSee systems, Inc. Lebanon, NH, U.S.A.), with the equations (Eqs. 1 to 3) integrated as shown in Figure S 3**.** Ambient MP concentrations (CMP) were allowed to vary randomly following a Poisson distribution as were the data presented in Gewert et al. (2017), whereas the CR values were normally distributed with a mean and SD of 1041 L $h^{-1}$ and 27 $h^{-1}$, respectively, and GER values varied randomly between 0.05 and 0.26 $h^{-1}$ without any assumption regarding the distribution (Table S 3). The final value of each run was used to represent an individual in the population.

### SI Tables and Figures

**Table S 1.** FTIR classification results based on the comparison with a reference database (Primpke et al. 2018). A match to one or more equally plausible compounds was accepted at HQI ≥ 70%. Spectra with HQI-scores below this threshold were classified as unknown.

| Sample | Sample type | Classification | Group |
|---|---|---|---|
| 1 | transparent fiber | fur | natural |
| 2 | transparent fiber | polyvinyl | MP |
| 3 | blue fiber | plant fiber, cellulose, hydroxypropyl methyl cellulose, polyvinyl alcohol | MP |
| 4 | transparent fiber | algae | natural |
| 5 | transparent fiber | fur | natural |
| 6 | transparent fiber | algae | natural |
| 7 | red fiber | fur | natural |
| 8 | black fiber | plant fiber | natural |
| 9 | black fiber | plant fiber, cellulose | natural |
| 10 | blue fiber | plant fiber, viscose | MP |
| 11 | blue fiber | viscose, polyvinyl | MP |
| 12 | transparent fiber | chitin | natural |
| 13 | blue fiber | unknown | unknown |
| 14 | blue fiber | unknown | unknown |
| 15 | blue fiber | plant fiber, cellulose, hydroxypropyl methyl cellulose, polyvinyl alcohol | MP |
| 16 | red fiber | plant fiber, cellulose, hydroxypropyl methyl cellulose, rubber | MP |
| 17 | black fiber | unknown | unknown |
| 18 | white fragment | acrylonitrile butadiene styrene, alkyd varnish | MP |
| 19 | blue fiber | unknown | unknown |
| 20 | black fiber | unknown | unknown |
| 21 | red fiber | rubber, ethylene propylene, poly 1-butene isotactic | MP |
| 22 | brown fragment | polyamide, fur | natural |
| 23 | black fiber | rubber, poly 1-butene isotactic | MP |
| 24 | black fiber | rubber, polyethylene | MP |
| 25 | blue fiber | fur | natural |
| 26 | red fiber | unknown | unknown |

**Table S 2.** Descriptive statistics for the microplastics recovered from the gastrointestinal tract of Baltic Sea herring as well as gut fullness of the examined fish. For each basin, the number of differently colored fragments and fibers that were recovered from the fish are shown as well as MP frequency of occurrence (FO), median, range (min-max) and mean; moreover, the mean values were calculated for all fish and for the fish that contained putative MP in their GIT (i.e., excluding zero values). Gut fullness (GF) is presented as percent of fish with empty stomachs, median gut fullness and the corresponding inter-quartile range (IQR). Values in parentheses represent FTIR-validated MP data. Data are ordered north to south.

| | Fibers | | | | Fragments | | | MP total | | | | | GF | | |
|---|---|---|---|---|---|---|---|---|---|---|---|---|---|---|---|
| | Black/Brown | Red | Blue | Clear | Black/Brown | Red | Green | FO (%) | Median | Mean all samples | Mean of presences | Range (min-max) | % fish with empty GIT | Median | IQR |
| Bothnian Bay | 0-7 | 0-38 | 0-0 | 0-3 | 0-0 | 0-3 | 0-12 | 46.7 (40.0) | 0 | 4.1 (3.5) | 8.9 (3.9) | 0-40 (0-15) | 10 | 50 | 25-50 |
| Bothnian Sea | 0-8 | 0-18 | 0-0 | 0-5 | 0-7 | 0-0 | 0-2 | 30.0 (12.5) | 0 | 1.3 (1.0) | 4.2 (3.4) | 0-18 (0-7) | 7.5 | 25 | 25-50 |
| Northern Baltic Proper | 0-1 | 0-0 | 0-0 | 0-51 | 0-0 | 0-2 | 0-2 | 30.0 (20.0) | 0 | 6.7 (6.6) | 22.2 (12.8) | 0-51 (0-20) | 25 | 25 | 19-50 |
| Western Gotland Basin | 0-1 | 0-3 | 0-1 | 0-1 | 0-0 | 0-7 | 0-0 | 40.0 (25.0) | 0 | 1.0 (0.9) | 2.5 (1.4) | 0-7 (0-3) | 10 | 25 | 25-25 |
| Bornholm Basin | 0-1 | 0-13 | 0-0 | 0-2 | 0-0 | 0-0 | 0-0 | 20.0 (15.0) | 0 | 0.9 (0.9) | 4.5 (2.3) | 0-13 (0-5) | 0 | 50 | 25-56 |
| **Total** | **0-8** | **0-38** | **0-1** | **0-51** | **0-7** | **0-7** | **0-12** | **33.8 (22.3)** | **0** | **2.7 (1.0)** | **7.8 (4.4)** | **0-51 (0-20)** | **9.2** | **25** | **25-50** |

**Table S 3.** Variables and simulation settings used to model microplastic ingestion in Baltic Sea herring. Details regarding derivation of the values are provided in the Supporting Information 2.1.

| Parameter | Unit | Average | Min | Max | S.D | Distribution | Species | Meaning | Reference |
|---|---|---|---|---|---|---|---|---|---|
| CMP | MP $L^{-1}$ | $5.8 \times 10^{-4}$ | | | | Poisson | | MP concentration in the water column | Gewert et al. 2017 |
| CR | L $ind.^{-1}$ $h^{-1}$ | $1.04 \times 10^{3}$ | | | $2.6 \times 10^{2}$ | Normal | *Clupea harengus* | Clearance rate | Varpe & Fiksen 2010 |
| GER[1] | $h^{-1}$ | | $5 \times 10^{-2}$ | $2.6 \times 10^{-1}$ | | | *Sardinops sagax*, *Sardina pilchardus* | Gut evacuation rate | Van der Lingen 1998, Costalago & Palomera 2014 |
| IR | MP $h^{-1}$ | | | | | | | Number of MP ingested at time *t* | |
| MP | MP | | | | | | | Number of MP in fish stomach at time *t* | |
| Eg | MP $h^{-1}$ | | | | | | | Number of MP egested at time *t* | |

1. The lower value for GER is based on data for *Sardinops sagax* (Van der Lingen 1998) while the higher is derived from *Sardina pilchardus* (Costalago & Palomera 2014).

**Table S 4.** Descriptive statistics for the predicted (modelled) and observed distributions of the MP burden in the Baltic herring. The data are presented as either "Total", i.e., where individuals without MP in the GIT are included, or "Zeros excluded" that shows only the fish with positive MP burden.

| | Total | | Zeros excluded | |
|---|---|---|---|---|
| | Mod | Obs | Mod | Obs |
| n | 1000 | 130 | 806 | 25 |
| Mean | 4.7 | 0.9 | 5.9 | 4.7 |
| SD | 4.7 | 3.0 | 4.5 | 5.6 |
| Median | 3.6 | 0.0 | 4.4 | 2.0 |
| Min | 0.0 | 0.0 | 1.3 | 1.0 |
| Max | 33.3 | 20.0 | 33.3 | 20.0 |
| Range | 33.3 | 20.0 | 32.0 | 19.0 |
| Skew | 1.8 | 4.4 | 2.0 | 1.5 |
| Kurtosis | 5.0 | 20.1 | 5.7 | 0.8 |
| SE | 0.1 | 0.3 | 0.2 | 0.1 |

**Table S 5.** Summary statistics and factor loadings for the variables used in the factor analysis. WS MP burden = weight specific MP burden. Factor loadings > 0.7 are considered statistically significant.

| | **Factor 1** | **Factor 2** |
|---|---|---|
| WS MP burden | -0.135 | 0.578 |
| BDE sum | **0.921** | 0.243 |
| HBCD | **0.997** | -0.042 |
| DD sum | **0.953** | 0.083 |
| HCB | **0.941** | -0.099 |
| PCB sum | 0.562 | **0.824** |
| SS | 3.969 | 1.091 |
| Proportion var | 0.662 | 0.182 |
| Cumulative var | 0.662 | 0.843 |

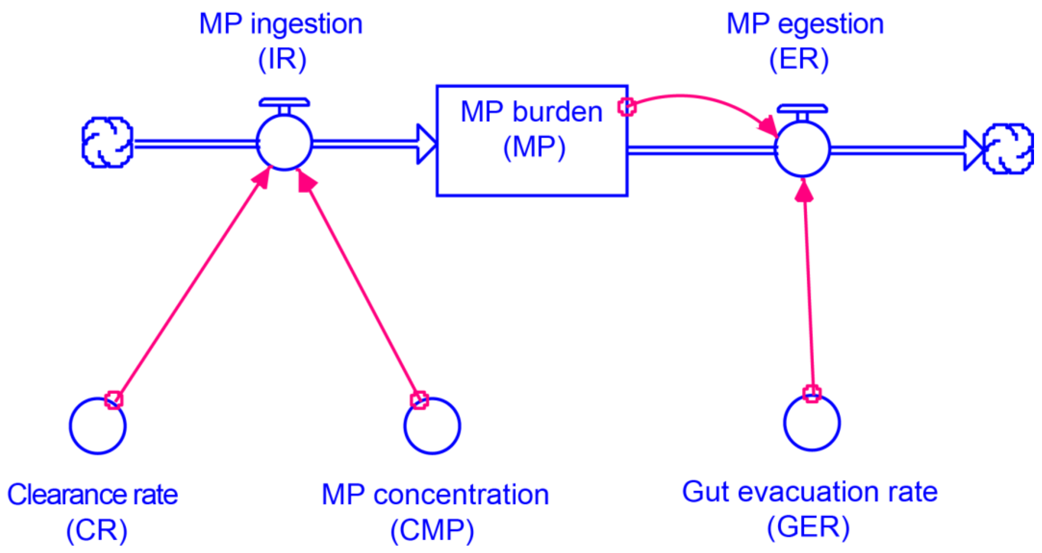

**Figure S 1.** Schematic representation of the model used to predict microplastic ingestion in Baltic herring.

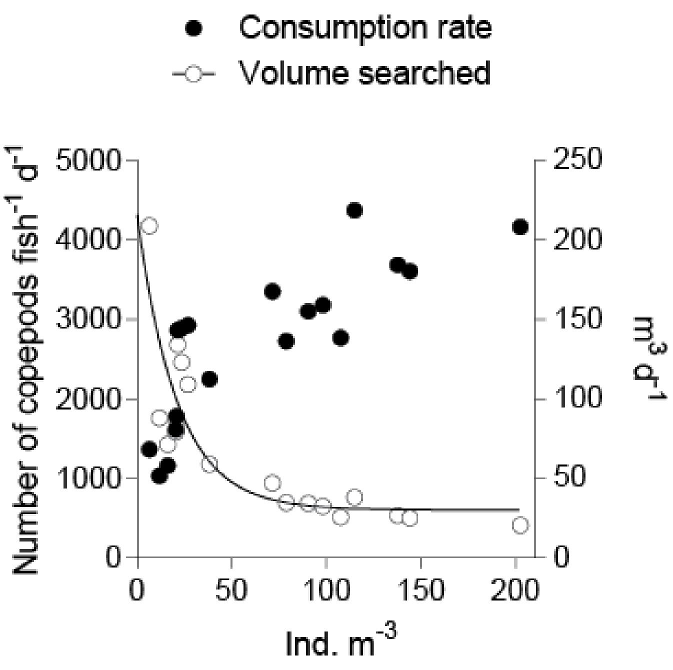

**Figure S 2.** Consumption rates (left axis) and clearance rates (right axis) as a function of *Calanus finmarchicus* abundance. The values are based on the data presented in Fig. 5, Varpe and Fiksen (2010) and adjusted for fish with average body weight of 35 g.

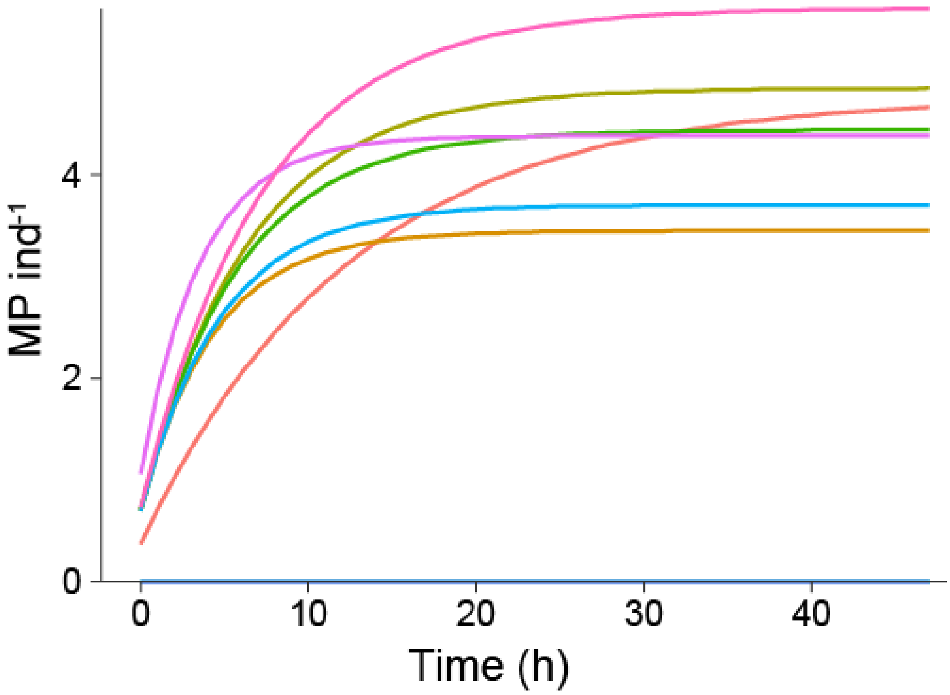

**Figure S 3.** Modeled MP burden (MP ind$^{-1}$) in the first ten simulation runs for 48 h. Observe that values are stabilized at the end of the simulation; these values are used to represent intrapopulation variability. Three out of ten individuals contain no MP.
